# From Eggs to Guts: Symbiotic Association of *Sodalis nezarae* sp. nov. with the Southern Green Shield Bug *Nezara viridula*

**DOI:** 10.1101/2024.10.18.619066

**Authors:** Magda A. Rogowska-van der Molen, Alejandro Manzano-Marín, Jelle L. Postma, Silvia Coolen, Theo van Alen, Robert S. Jansen, Cornelia U. Welte

## Abstract

Phytophagous insects engage in symbiotic relationships with bacteria that contribute to digestion, nutrient supplementation, and development of the host. The analysis of shield bug microbiomes has been mainly focused on the gut intestinal tract predominantly colonized by *Pantoea* symbionts, and other microbial community members in the gut or other organs have hardly been investigated. In this study, we reveal that the Southern green shield bug *Nezara viridula* harbours a *Sodalis* symbiont in several organs, with a notable prevalence in salivary glands, and anterior regions of the midgut. Removing external egg microbiota via sterilization profoundly impacted insect viability but did not disrupt the vertical transmission of *Sodalis* and *Pantoea* symbionts. Based on the dominance of *Sodalis* in testes, we deduce that *N. viridula* males could be involved in symbiont vertical transmission. Genomic analyses comparing *Sodalis* species revealed that *Sodalis* sp. Nvir shares characteristics with both free-living and obligate insect-associated *Sodalis* spp. *Sodalis* sp. Nvir also displays genome instability typical of endosymbiont lineages, which suggests ongoing speciation to an obligate endosymbiont. Together, our study reveals that shield bugs harbour unrecognized symbionts that might be paternally transmitted.

## Introduction

Insects establish symbiotic relationships with microorganisms which can confer important physiological traits to their host. In this way, microbes allow insects to adopt new lifestyles which may enable them to colonize diverse plant species (Dillon & Dillon, 2004; van den Bosch & Welte, 2017). Particularly plant-feeding true bugs (Hemiptera) largely rely on their symbionts (Colman et al., 2012; Feng et al., 2019; Kikuchi, Hayatsu, et al., 2012; Salem et al., 2014). Within hemipterans, Pentatomidae shield bugs represent the largest family, including 8 000 species of which several are crop pests (McPherson et al., 2017). One such insect is *Nezara viridula*, also known as the Southern green shield bug, a polyphagous species that feeds on over 30 plant families including crops, thereby causing major economic losses worldwide (McPherson & McPherson, 2000). Current pest control methods rely on insecticides, even though they were found to be ineffective against shield bugs. Recent studies, however, highlighted the potential of targeting symbiotic microorganisms as an alternative pest control strategy (Chung et al., 2018; Gonella & Alma, 2023). *N. viridula* microbiota has been shown to aid insects in overcoming plant defences by deactivating protease inhibitors and degrading soybean isoflavonoids, and the leguminous toxin 3-nitropropionic acid (Medina et al., 2018; Rogowska-van der Molen et al., 2022; Zavala et al., 2015). Although *N. viridula* microbiota has been studied and is known to be beneficial to the host (Medina et al., 2018; Prado et al., 2009; Tada et al., 2011), the composition and function of microbiota on the surface of eggs has until recently remained poorly characterized (Geerinck et al., 2022). A better understanding of the host-microbe relationships in *N. viridula* could eventually contribute to the development of targeted pest control (Gonella & Alma, 2023; Rogowska-van der Molen et al., 2023).

Many *N. viridula* symbionts are essential for the viability of their hosts. For example, experimental removal of the *N. viridula* egg microbiota with surface sterilization disrupted nymphal colonization by symbionts and severely increased nymphal mortality (Tada et al., 2011). This underscores the critical nature of symbionts for the development and survival of insects. Moreover, several studies have described the co-evolution of hemipterans with their symbionts by the development of specialized organs to house and sustain a stable symbiotic population. In Pentatomidae, obligate *Pantoea* symbionts are typically harboured in crypts of the posterior M4 region of the gut (Bistolas et al., 2014; Duron & Noel, 2016; Hosokawa et al., 2016; Kikuchi, Hosokawa, et al., 2012; Matsuura et al., 2012; Taylor et al., 2014). Next to *Pantoea* located in crypts, it has been recently suggested that shield bugs harbour a *Sodalis* strain as a core symbiont in anterior gut compartments that likely contributes to thiamine supplementation (Fourie et al., 2023). Several studies have reported the occurrence of *Sodalis* in various insects and characterized them as obligate or facultative symbionts. An obligate *Sodalis* symbiont of slender pigeon louse *Columbicola columbae* was found to be maternally transmitted in bacteriocytes, supporting their host insect by supplementing nutrients and participating in digestion (Fukatsu et al., 2007). Moreover, several co-obligate *Sodalis* symbionts have been identified in aphids (Manzano-Marín et al., 2023; Manzano-Marin et al., 2017), and Garber et al. (2021) described the recent acquisition of intracellular *Sodalis* endosymbionts in mealybugs. Also, ’*Candidatus* Sodalis baculum’, an intracellular symbiont residing in the bacteriome of the seed bug *Henestaris halophilus* (Heteroptera) was reported to complement the host’s diet (Santos-Garcia et al., 2017). On the other hand, *Sodalis*- allied bacteria can function as facultative symbionts in acorn weevils and psyllids, although their symbiotic role remains unclear (Ghosh et al., 2020; Kaiwa et al., 2010; Toju & Fukatsu, 2011). Regarding shield bugs, while some authors have suggested a secondary role of *Sodalis* (Hosokawa et al., 2015), a recent study showed a high abundance of both *Sodalis* and the obligate *Pantoea* symbiont in the egg microbiome, suggesting a similar vertical transmission route of both symbionts (Geerinck et al., 2022).

As most research on shield bug symbionts has focused on the obligate *Pantoea* <u>symbionts</u> and on the M4 crypts, other bacterial species like *Sodalis* have been hardly investigated. Consequently, we currently lack information about the abundance of *N. viridula-*associated bacteria in organs other than the gut. In this study, we aimed at elucidating the transmission mechanisms and localisation of the suspected main core symbionts (*Pantoea* and *Sodalis*) in *N. viridula*. First, through egg surface sterilisation, we removed the symbionts and used 16S rRNA gene profiling to determine the transmission of *Sodalis* and *Pantoea*. The presence of bacteria in *N. viridula* organs was then visualized with confocal microscopy with a focus on *Sodalis*. The results indicated a high prevalence of the symbiont in salivary glands, testes, and anterior regions of the midgut. Finally, a comparative genomic analysis unveiled the divergent nature of the *Sodalis* symbiont of *N. viridula* (hereafter *Sodalis* sp. Nvir) which displays typical endosymbiotic lifestyle traits which result from long-term vertical transmission. These findings provide valuable insights into understanding the symbiont transmission route in pest shield bugs and broaden our knowledge of *Sodalis* spp. as insect-associated bacteria.

## Materials and Methods

### Insect collection and rearing

*Nezara viridula* shield bugs were collected in the field on creeping thistle (*Cirsium arvense*) in the Netherlands (51.348028, 6.128802) in July 2019. Given the susceptibility of *N. viridula* for inbreeding, adult insects were regularly introduced to the established population. Insects were introduced in March 2021 from the local populations in the Netherlands (De Lier, Rotterdam, and Bleiswijk), in March 2023 from Wageningen Plant Research (Bleiswijk, the Netherlands) and in April 2023 from Koppert (Berkel en Rodenrijs, the Netherlands). The insects were transferred to a greenhouse and placed in a rearing cage (90x60x60 cm) to establish a colony. *N. viridula* individuals were reared in a greenhouse facility with no humidity control at room temperature with normal daylight and additional light to obtain a photoperiod of 16:8 h (light:dark) year-round. Insects were provided with sunflower (*Helianthus annuus*), soybean (*Glycine max*), brown mustard (*Brassica juncea*) seeds, flat beans (*Phaseolus vulgaris*), and the native plants crown vetch (*Securigera varia*), black mustard (*Brassica nigra*) and black nightshade (*Solanum nigrum*).

### Egg sterilization

Six one-day old laid egg masses were collected from the *N. viridula* population. Three masses (63, 75, 76 eggs) were surface sterilized by submerging in 1 mL 70% ethanol for 1 min. As a control, three egg masses (59, 65, 78) were treated with 1 mL autoclaved demineralized water for 1 min. For detection of *Sodalis* and *Pantoea* symbionts in and outside of eggs, one sterilized (48 eggs) and one control egg mass (47 eggs) was crushed using a sterile pestle after which DNA was isolated and used for diagnostic PCR.

To assess the influence of microbiota removal on the survival and composition of *N. viridula* microbiota, two sterilized (47, 48 eggs) and two control egg masses (42, 47 eggs) were placed in separate rearing cages at room temperature containing sunflower, soybean, brown mustard seeds and flat beans. The survival rate was monitored by tracking the number of insects until reaching adulthood. To analyse the gut microbial community, two female adult and one male adult insects were randomly selected from each cage. The individuals were dissected, and DNA was extracted from their gut systems. The gut microbiota composition was determined using 16S rRNA gene amplicon sequencing.

### Insect dissection

To determine the effect of surface sterilization on microbiota composition, complete gut systems (M1- M4 midgut and hindgut) of adult *N. viridula* were dissected directly after submersion of insects in 70% ethanol for one minute. Dissection was performed under non-sterile conditions using a stereomicroscope, scalpel, and forceps. Separation of tissues from the insect body was performed in phosphate-saline buffer (PBS; 137 mM NaCl, 2.7 mM KCl, 10 mM Na2HPO4 and 1.8 mM KH2PO4, pH 7.4) to prevent tissue rupture. Gut systems were disrupted vortexing in 100 µL PBS for DNA isolation.

### DNA isolation

Isolation of the DNA for 16S rRNA gene amplicon sequencing and diagnostic PCR was performed with a DNeasy PowerSoil kit (QIAGEN, the Netherlands). Disrupted tissue, eggs and plant pellets were transferred into the lysis buffer in the Powerbead tubes and vortexed (10 min, 50 Hz) with TissueLyser LT (QIAGEN). DNA was eluted in 40 µL Nuclease Free Water (Thermo Fisher Scientific Inc., Waltham, USA) and quantified with a Qubit dsDNA HS assay kit (Thermo Fisher Scientific Inc.). Before PCR, DNA concentrations of organ samples had been standardized to the equal concentration of 2 ng µL^-1^.

### 16S rRNA gene amplicon sequencing and analysis

The gut bacterial community of *N. viridula* was determined by amplification of the V3-V4 region of the 16S rRNA gene. The sequencing was performed by Macrogen (the Netherlands) with *Bac341F* and *Bac806R* primers (Caporaso et al., 2012; Herlemann et al., 2011) using an Illumina MiSeq sequencer (Illumina). Paired-end (2x 301bp) reads libraries were prepared with the Herculase II Fusion DNA Polymerase Nextera XT Index Kit V2 (Illumina).

The quality of the raw paired-end sequences was checked with FastQC 0.11.8 (Bushnell, 2014). The reads were then filtered, and adapters were trimmed. Approximately 40 000-70 000 paired-end sequencing reads were obtained per sample. The data were further processed using the DADA2 1.8 pipeline (Callahan et al., 2016) in R. Phylogenetic taxonomy of the reads was assigned using the SILVA 16S rRNA gene database 138.1 (Quast et al., 2012). Count data were normalized to relative abundance. Microbial community profiles were analysed and amplicon sequence variants (ASVs) were visualized using the phyloseq (McMurdie & Holmes, 2013) and ggplot2 (Wickham, 2009).

### Genomic analysis

The previously published genome of *Sodalis* sp. Nvir (Coolen et al., 2024) was further analyzed in this study. A draft annotation was performed using Prokka v1.4.6 (Supplementary Information Methods) (Seemann, 2014). To infer pseudogenes, PseudoFinder v1.1.0 (Syberg-Olsen et al., 2022) was run with *S. praecaptivus* HS1 reference and without dN/dS analysis. To analyse rearrangements among strains related to HS1, OrthoVenn3 and OrthoFinder were used (e-value 1x10^-3^; 1.5 inflation value) (Emms & Kelly, 2019; Sun et al., 2023).

For phylogenetic placement of the previously isolated *Sodalis* sp. Nvir (Coolen et al., 2024), we collected genomes of a comprehensive dataset of *Pectobacteriaceae*, “*Bruguierivoracaceae*” (Li et al., 2021) (Supplementary Information Methods). Phylogenetic inference was done using IQtree v2.2.2.7 (LG4X+I+G; (Minh et al., 2020)) with 1000 UltraFast bootstrap replicates (Hoang et al., 2018). The resulting tree was visualised and exported for editing using Figtree v1.4.4 (https://github.com/rambaut/figtree). Additionally, ANI, 16S rRNA gene identity and phage regions were identified (Supplementary Information Methods).

### Diagnostic PCR

The presence of *Sodalis* and *Pantoea* symbionts in sterilized and control egg masses and *Sodalis* in organs of adult *N. viridula* male and female was determined with a PCR reaction quantifying *gro*L gene using specific primers (Supplementary Information Methods). As a positive control, DNA from *Sodalis*sp. Nvir(Coolen et al., 2024) was used, whereas Nuclease Free Water (Thermo Fisher Scientific Inc.) served as a negative control. Amplification was performed using SensoQuest lab-cycler (BIOKÉ, the Netherlands) in a 25 mL reaction volume (Supplementary Information Methods). PCR products were examined by electrophoresis under UV light.

### Fluorescence *in situ* hybridization

To image total bacteria, Gammaproteobacteria and *Sodalis*, we dissected the fat body, ovary, testes, salivary glands, M1, M2, M3, M4 sections of the midgut, hindgut, and Malpighian tubules from three adult males and three adult *N. viridula* females. With images collected from individual males and females, one representative image is shown here.. Samples were hybridized with a Fluos-labelled general bacterial probe (Eub-mix; (Amann et al., 1990; Daims et al., 1999)) and a Cy5-labeled Gammaproteobacterial probe (GAM42A; (Manz et al., 1992)) along with a GAM42A competitor probe were employed. A Cy3-labeled probe specific for *Sodalis* sp. was used for targeted detection (Sod1238R; (Koga et al., 2013)). Samples were visualized using laser scanning microscopy. For a detailed protocol see Supplementary Information Methods.

## Results

### Egg-surface sterilization decreases *N. viridula* survival

Symbionts are instrumental to the development and fitness of *N. viridula.* However, upon the recent discovery of an internal egg microbiota (Geerinck et al., 2022) we questioned how the removal of symbionts from the egg surface impacts *N. viridula* survival rate as well as symbiont inheritance (Figure 1). We observed that upon symbiont removal from the egg surface, the survival rate of *N. viridula* decreased to 11% while the control population reached 86% (Figure 1A). Moreover, the removal of symbionts negatively affected the appearance of adult females (Figure 1B) and resulted in discolouration and retarded growth. To assess the effect on the microbiome, we applied 16S rRNA gene amplicon sequencing, which revealed that the gut microbial community of control non-sterilized insects was largely dominated by *Pantoea* and *Sodalis* (Figure 1C). These genera were also dominant in sterilized insects, but the relative abundance of *Sodalis* was significantly lower in sterilized insects compared to the control population (45% vs 20%, Student’s *t-*test; *P*-value <0.05). In addition, in two sterilized insects, the relative abundance of *Sodalis* decreased in favour of *Enterobacter* and *Enterococcus*, suggesting that egg-surface sterilization allowed for colonization of the gut by bacteria from the surrounding environment. Altogether, these results demonstrate that the removal of symbionts from the egg surface affects insect survival and external appearance, while the two main symbionts *Pantoea* and *Sodalis* still dominate the gut microbial community.

**Figure 1.**
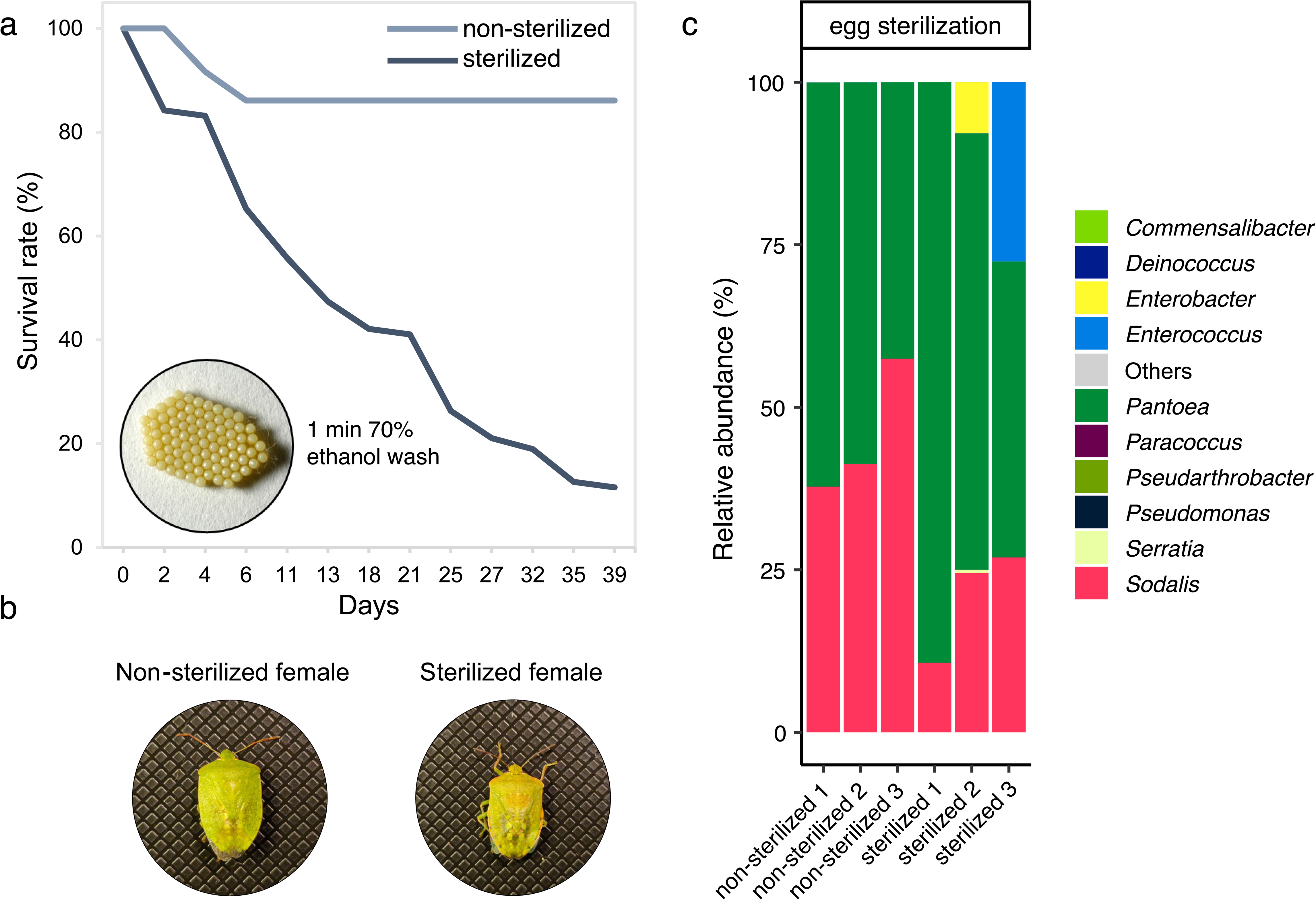
The effect of egg-surface sterilization on survival and gut microbial community of *Nezara viridula*. **A)** Survival rate was monitored from hatching until reaching adulthood in non-sterilized (control) and egg-surface sterilized populations of *N. viridula*. Two egg clusters in a sterilized population (n = 95)were treated with a 1 min wash in 70% ethanol while two non-sterilized egg clusters (n = 89) were subjected to 1 min wash in demineralized autoclaved water. **B)** External appearance of the control and treated adult females. **C)** Gut microbial community composition in adult *N. viridula* individuals based on 16S rRNA gene amplicon sequencing subjected to symbiont removal via egg-surface sterilization or non-sterilized. The taxonomy is displayed at the genus level. ‘Others’ represent the amplicon sequence variants (ASVs) that average below 0.5% of all reads. Individual bar graphs represent the sequencing of one adult insect per bar, with three biological replicates (n = 3) per treatment group.

### *Sodalis* dominates in salivary glands and the anterior section of the midgut

*N. viridula* and other Pentatomidae insects are commonly associated with obligate *Pantoea* symbionts, located in the M4 crypts, however little is known regarding their other symbiotic partners. Here, we investigated the prevalence of the symbiont *Sodalis* in organs and eggs. Hatching shield bug nymphs acquire symbionts that colonize the gut tract via ingestion of maternally smeared symbionts from the egg surface (Figure 2A) (Hosokawa et al., 2016), so surface-sterilization of egg clusters probably disturbs their acquisition. We compared surface-sterilized to surface-washed egg clusters of *N. viridula* using diagnostic PCR; this confirmed the presence of *Pantoea* and *Sodalis* in non-sterilized eggs, and also detected DNA of *Pantoea* and *Sodalis* symbionts in surface-sterilized egg cluster, corroborating the recently reported internal egg microbiome (Geerinck et al., 2022).

**Figure 2.**
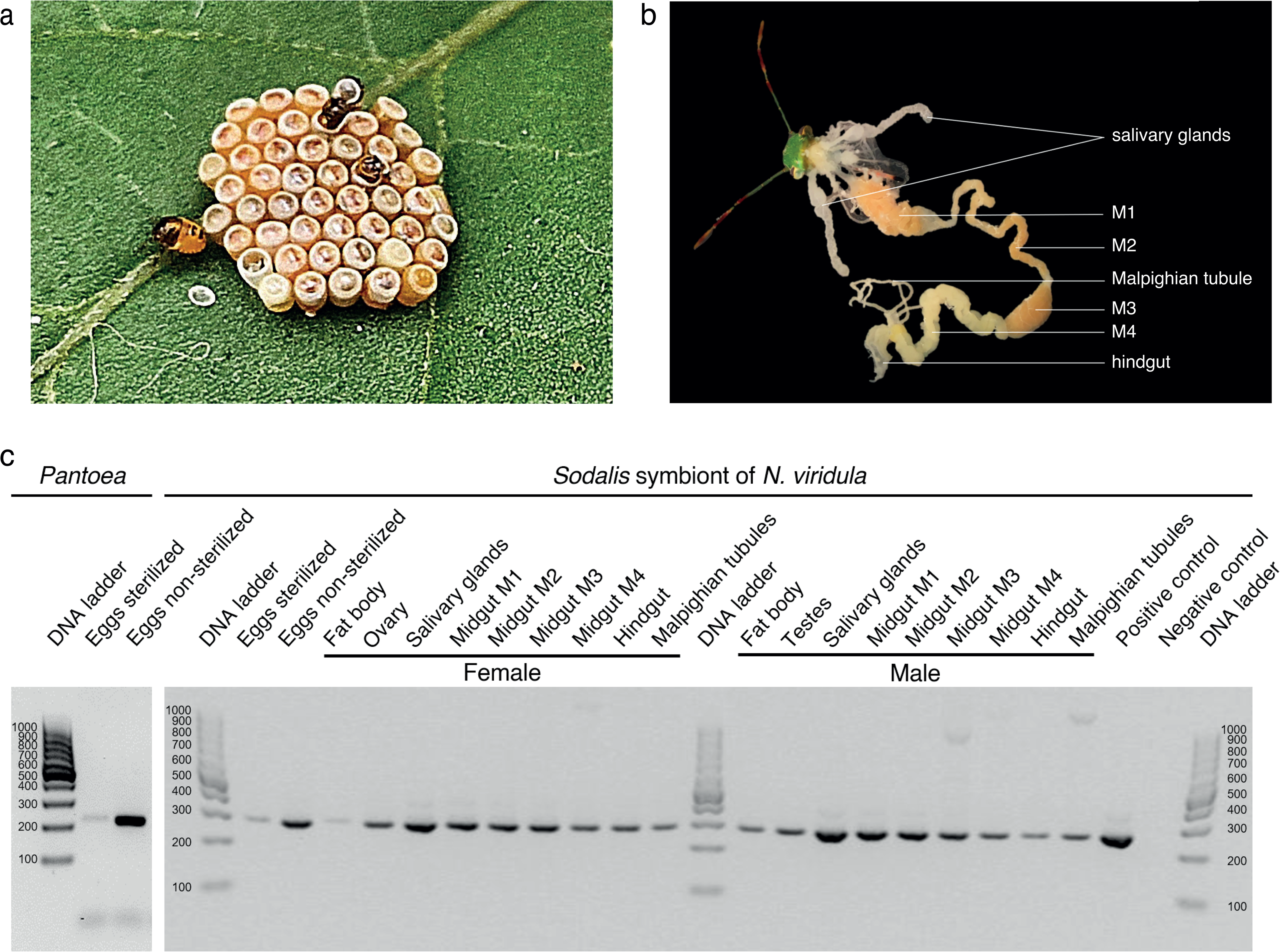
*Sodalis* distribution in *Nezara viridula*. **A)** Hatching of nymphs from the egg of *N. viridula*. **B)** Organization of organs dissected from adult *N. viridula*. M1, midgut first section; M2, midgut second section; M3, midgut third section; M4, midgut fourth section with crypts. **C)** Diagnostic PCR detection of obligate *Pantoea* and *Sodalis* symbionts in non-sterilized (n = 47) and surface-sterilized eggs (n = 48) and *Sodalis* in organs of female and male adult *N. viridula*. Negative control was nuclease-free water while positive control was the DNA extracted from the isolated *Sodalis* strain from the *N. viridula* salivary glands. DNA extracted from eggs, organs and tissue was normalized to an equal concentration of 2 ng µL^-1^.

Next, we sought to investigate the localization of the *Sodalis* symbiont in organs of *N. viridula* (Figure 2B-C). Diagnostic PCR analysis of dissected tissues detected the symbiont DNA in all *N. viridula* organs with the most prominent band in salivary glands. Notably, *Sodalis* was present throughout the entire gut system and it was observed in reproductive organs, as well as in the Malpighian tubules.

To verify the localization of *N. viridula*-associated microbiota and the *Sodalis* symbiont, we performed fluorescence *in situ* hybridization (FISH) on organs and visualized them with confocal laser scanning microscopy. Our analysis revealed that bacteria formed clusters in the entire principal salivary gland (Figure 3A-B). The tissue was colonized with Gammaproteobacteria, of which *Sodalis* was a dominant member. The gut of shield bugs is a complex alimentary system colonized with symbionts and is divided into a midgut that consists of four sections (M1-M4), and a hindgut (Figure 2B) (Tada et al., 2011). Confocal microscopy of the gut system indicated the presence of primarily Gammaproteobacteria in all gut sections, and *Sodalis* in M1 and M3 (Figure 4). In addition, M1 seemed to harbour more *Sodalis* than M3, whereas no *Sodalis* was found in the M2 and M4 sections or the hindgut. The M4 section was densely colonized with Gammaproteobacteria other than *Sodalis* which were largely localised to the surrounding crypts (Figure 4D). Together, the PCR and FISH imaging observations demonstrate that the *Sodalis* symbiont displays restricted localisation in several *N. viridula* digestive organs.

**Figure 3.**
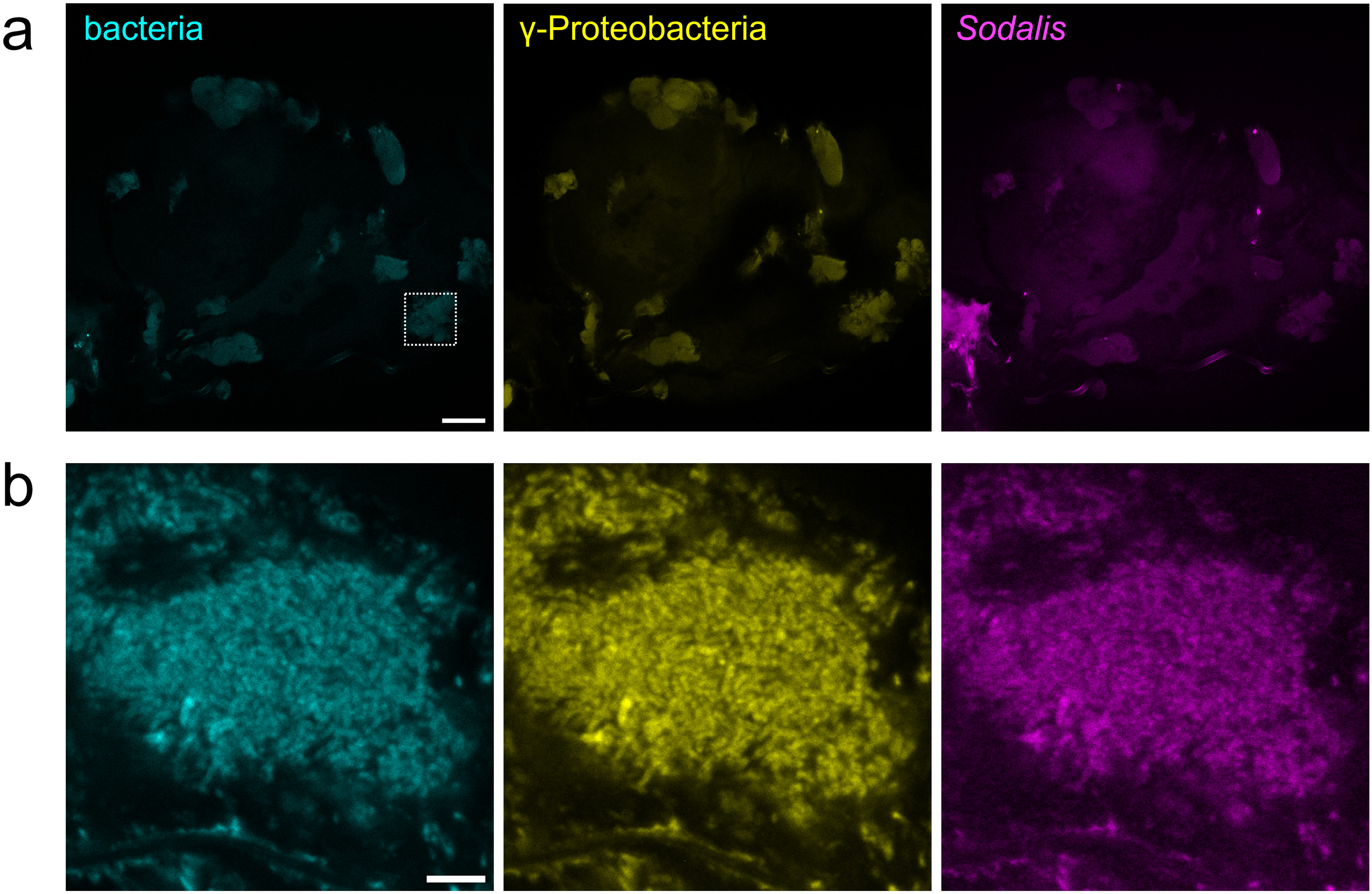
Colonization of *Nezara viridula* salivary glands by Gammaproteobacteria and *Sodalis*. Confocal laser scanning micrographs show the adult *N. viridula* salivary gland where bacteria are visualized with fluorescence *in situ* hybridization. **A)** Overview micrographs of a principal salivary gland. A dashed square in the left panel marks the localization of the magnified panel in B) Scale bar = 100 µm. Left panel, detection of all bacteria with probe EUB-mix in cyan (Fluos); middle panel, γ- proteobacteria detected with probe GAM42A in yellow (Cy5); right panel, *Sodalis* detected with probe Sod1238R in magenta (Cy3) **B)** Magnified confocal micrographs indicated in panel A) with a dashed square of the principal salivary gland. Scale bar = 5 µm. The images shown here are representative of FISH micrographs collected from three individual insects (n=3).

**Figure 4.**
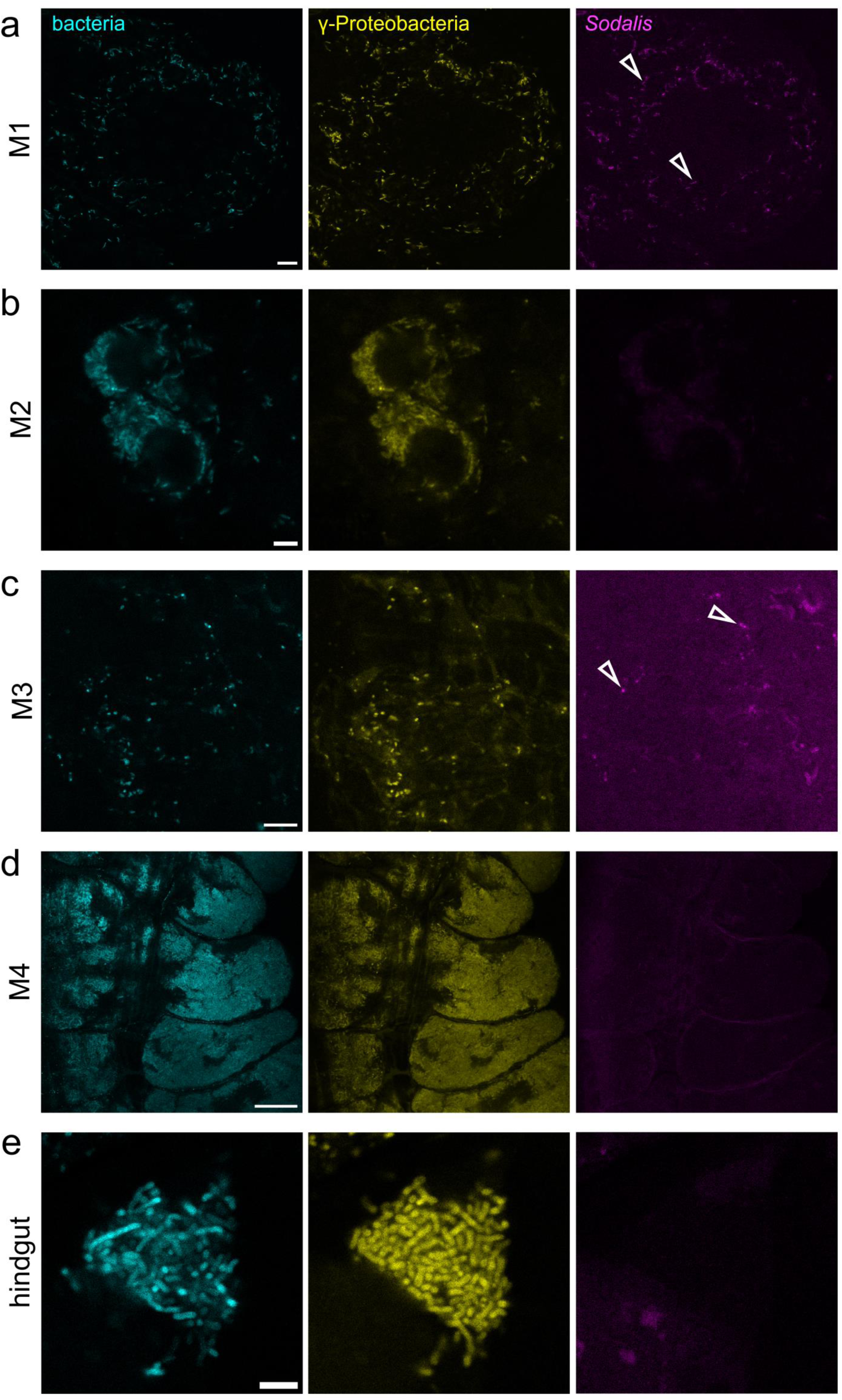
Colonization of the *Nezara viridula* gut by Gammaproteobacteria and *Sodalis*. Confocal micrographs show the adult *N. viridula* intestinal tract. Panel *bacteria* shows FISH-probe *Eub-mix* in cyan (Fluos). Panel *γ-Proteobacteria* shows FISH-probe *GAM42A* in yellow (Cy5). Panel *Sodalis* shows FISH-probe *Sod1238R* in magenta (Cy3). Open arrowhead points at FISH-stained *Sodalis*. **A)** M1 section of the gut. Scale bar = 10 µm. **B)** M2 section of the gut. Scale bar = 10 µm. **C)** M3 section of the gut. Scale bar = 10 µm. **D)** M4 section of the gut with crypts. Scale bar = 25 µm. **E)** Hindgut. Scale bar = 5 µm. The images shown here are representative of multiple FISH micrographs collected from three individual insects (n=3).

### The presence of *Sodalis* in testes suggests paternal symbiont transmission

According to the diagnostic PCR analysis, *Sodalis* DNA was present in reproductive organs and Malpighian tubules. Thus, to confirm these results, we performed confocal microscopy of the reproductive organs. FISH microscopy indicated a lack of bacteria in the ovary (Supplementary Figure 1A). This finding corroborates the recently reported absence of microbes from dissected ovaries and unlaid eggs (Geerinck et al., 2022; Kikuchi et al., 2009). Nonetheless, our results revealed a dense population of Gammaproteobacteria and *Sodalis* in the testes (Figure 5A) suggesting a possible paternal involvement in symbiont transmission. Besides this, Malpighian tubules were colonized with Gammaproteobacteria (Figure 5B), and only a few microbes were seen in the fat body (Supplementary Figure 1B). Since those microbes were localized on the tissue surface, they were unlikely to be part of the fat body microbiome and rather represent bacteria that were part of the gut microbiome. Taken together, we show the symbiont distribution in *N. viridula* organs and provide the first evidence of the possible participation of shield bug males in the vertical transmission of symbionts.

**Figure 5.**
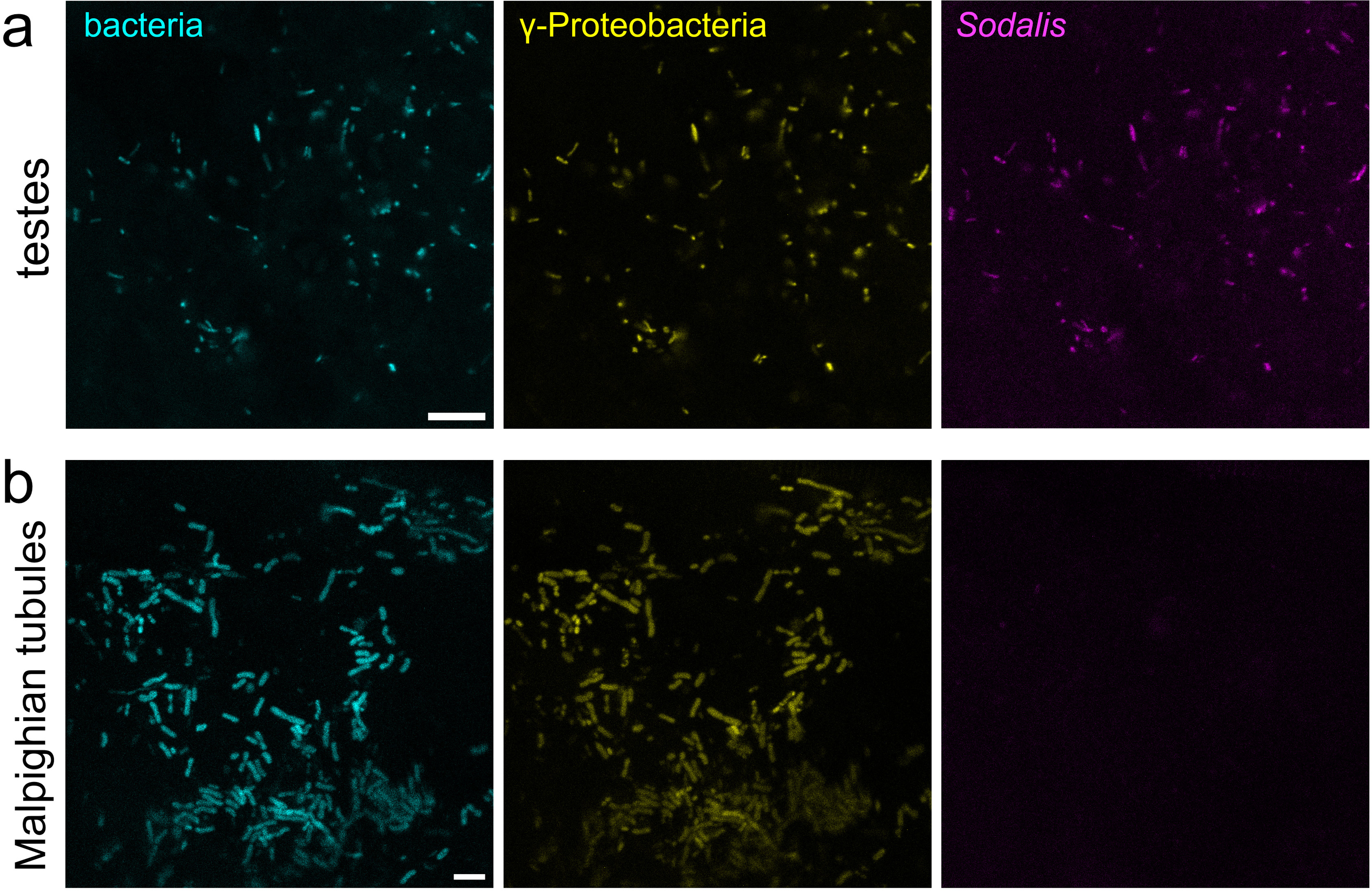
**Colonization of *Nezara viridula* testes and Malpighian tubules by Gammaproteobacteria and *Sodalis.*** Confocal micrographs show the adult *N. viridula* testes A) and B) Malpighian tubules. Panel *bacteria* shows FISH-probe *Eub-mix* in cyan (Fluos). Panel *γ-Proteobacteria* shows FISH-probe *GAM42A* in yellow (Cy5). Panel *Sodalis* shows FISH-probe *Sod1238R* in magenta (Cy3). **A)** Testes. Scale bar = 10 µm. **B)** Malpighian tubules. Scale bar = 5 µm. The images shown here are representative of multiple FISH micrographs collected from three individual insects (n=3).

### Phylogenetic placement and genomic characteristics indicate that *Sodalis* sp. Nvir is a novel beneficial symbiont

Due to the non-universal presence of *Sodalis* within other shield bug species, several authors have suggested a facultative role of *Sodalis* symbionts associated with this group of insect species (Kaiwa et al., 2010, 2011; Matsuura et al., 2014). However, the high abundance of *Sodalis* sp. Nvir in specific digestive organs of *N. viridula*, as well as the shared vertical transmission route with the obligate *Pantoea* symbionts, made us further questions the nature of this association (Ferrari & Vavre, 2011). To unveil the role of *Sodalis* sp. Nvir we characterized its genome and placed the strain in a phylogenetic tree including non-host associated and endosymbiotic *Sodalis* strains (Figure 6A and Supplementary Figure S2). The 16S rRNA gene identity values of *Sodalis* sp. Nvir and its close relatives suggested that *S. praecaptivus*, *S. pierantonii*, *S. melophagi*, and strains TME1 and Nvir all belonged to the same 16S- rRNA gene defined molecular species (Figure 6A and Supplementary Table S1; (Kim et al., 2014)). Moreover, the comparison of *Sodalis* sp. Nvir with its closest relatives suggested that Nvir finds itself in an intermediate genome degeneration stage between *S. praecaptivus* and *S. pierantonii* strain SOPE, implying Nvir’s recent transition to a vertically transmitted symbiotic lifestyle (Figure 6B).

**Figure 6.**
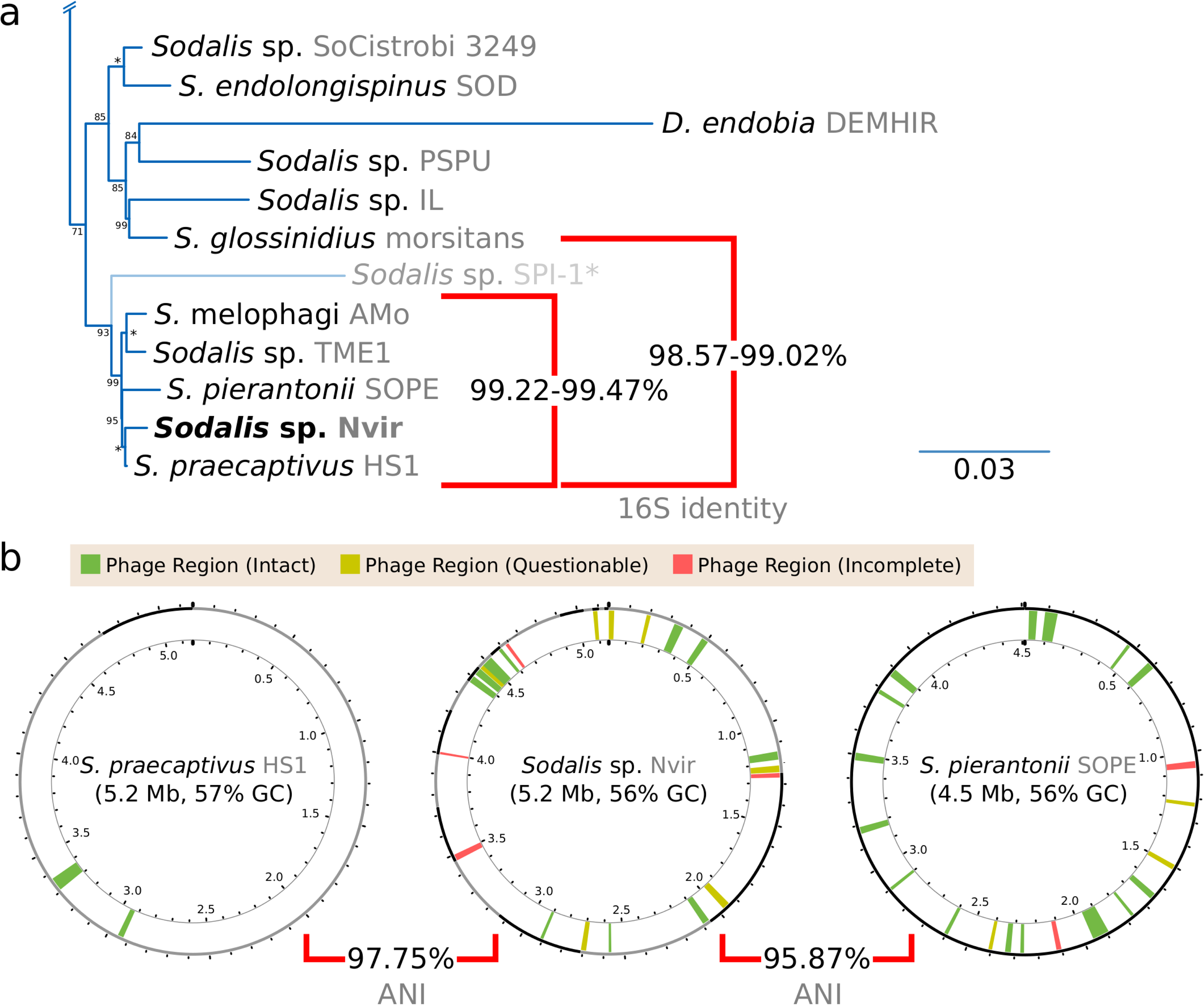
Phylogenetic placement and genomic features of *Sodalis* sp. Nvir. **(A)** Excerpt of Maximum-Likelihood phylogenetic placement of the novel *Sodalis* sp. Nvir within the *Sodalis* clade. Names on leaves specify the bacterial genera, species, and strains (in grey). Red squared brackets denote a two-way comparison of 16S rRNA gene identity values. Values at nodes indicate the *UltraFast* bootstrap support values in percentages. An asterisk (*) denotes a support of 100%. *Sodalis* sp. SPI-1 was excluded for the visual grouping representing 16S rRNA gene identity value scores. **(B)** Circular diagrams of selected *Sodalis* genomes. Numbers at inner ticks denote the location within the genome in Mega base pairs. The outer ring represents contigs/molecules of the genome assembly, with alternate grey and black colours separating them. Phage regions are coloured as specified in the colour key at the top of the diagrams. Squared brackets indicate the two-way ANI values.

To gain insight into the genomic characteristics of the *Sodalis* sp. Nvir symbiont, we performed a draft genome annotation and compared general genomic characteristics with other *Sodalis* strains (Table 1). Strain Nvir shows a similar genome size and G+C content as HS1 but contrastingly codes for about 200 transposase genes of IS elements. In addition, the analysis suggested a large number of pseudogenes and plasmids as well as a reduction of tRNA genes when compared to HS1, thus revealing clear signatures of genome reduction (Campbell et al., 2017; Campbell et al., 2015; Van Leuven et al., 2014). Moreover, the analysis showed that Nvir preserves a more intact genome than SOPE, altogether suggesting a recent history of association with its host marked by repeated genetic bottlenecks, typical of vertically transmitted endosymbionts.

**Table 1.** Genomic features of selected *Sodalis* spp. General genomic features evidencing genome reduction in *Sodalis* spp.

| Strain | *S. praecaptivus* HS1 | *Sodalis* sp. Nvir | *S. pierantonii* SOPE |
| --- | --- | --- | --- |
| genome size | 5.2 Mbp | 5.2 Mbp | 4.5 Mbp |
| G+C content* | 57.47% | 56.09% | 56.06% |
| CDSs | 4,358 | 4,066 | 2,309 |
| pseudogenes | 75 | 1,108 | 1,771 |
| tRNAs | 76 | 66 | 55 |
| plasmids | 1 | 6 | none |

\*G+C content of the chromosome

To infer possible rearrangements among the closely related *S. praecaptivus* HS1, *S. pierantonii* SOPE, and *Sodalis* sp. Nvir, we predicted shared clusters of orthologous proteins (Supplementary Figure 3). By analysing the arrangement of these genes across strains, it became evident that strain Nvir has undergone widespread genome rearrangements, mirroring what is observed in the obligate endosymbiont *S. pierantonii* (Oakeson et al., 2014) and other symbiotic taxa (Manzano-Marin & Latorre, 2014), while contrasting what is observed in the genome of the facultative endosymbiont *S. glossinidius* (Clayton et al., 2012). In addition, the collinearity of the largest plasmid of strain Nvir (pl01) to the HS1’s plasmid indicated a common ancestor with pl01, despite the smaller size of Nvir’s plasmid (449 vs. 221 kbp).

Lastly, to predict possible nutritional contributions of the novel *Sodalis* sp. Nvir, we annotated the genes involved in the biosynthesis of essential amino acids, B vitamins, and other cofactors (Supplementary Tables S2-3). Compared to strain SOPE, Nvir retained a much larger metabolic repertoire related to the aforementioned compounds. However, the inactivation of certain pathways, when compared to HS1, was evidenced by the pseudogenisation and loss of genes. Unlike the obligate nutritional *Pantoea* symbiont of *N. viridula*, *Sodalis* sp. Nvir retains the ability to synthesise thiamine, an essential B vitamin. This metabolic difference resembles the proposed dual symbiotic system of the two-spotted stink bug *Bathycoelia distincta*, where the *Sodalis* symbiont genomes preserve the capacity for thiamine biosynthesis while the *P. bathycoeliae* symbiont has lost this capacity (Fourie et al., 2023). Lastly, while genes necessary for amino acid biosynthesis are preserved, Nvir displays loss of redundancy when compared to HS1.

In conclusion, our results suggest that divergent strain Nvir underwent speciation from its once free-living ancestor *S. praecaptivus*, as supported by the genome characterization, FISH imaging of *N. viridula* organs, and displayed vertical transmission route. Given the tradition of assigning different specific names to insect host-associated *Sodalis* endosymbionts that are vertically transmitted and beneficial to their hosts, we propose that *Sodalis* sp. Nvir represents a novel species within the *Sodalis* genus: ‘*Sodalis nezarae’* sp. nov., with strain Nvir being the sole sequenced representative of this taxon.

### Description of the species ‘*Sodalis nezarae*’

*Sodalis nezarae* (ne.za’rae. N.L. gen. n. *nezarae*, of the shield bug genus *Nezara*, in which the species exists as a vertically transmitted beneficial symbiont endosymbiont).

We propose the specific name *Sodalis nezarae* for the *Sodalis* endosymbiont of *N. viridula*. This symbiont was shown to colonize salivary glands, anterior regions of the midgut and testes, as confirmed by diagnostic PCR, using *Sodalis*-specific primers *SodF* (5’- CCCTTATCGATAGCCGCGTT-3’) and *SodR* (5’-GATCTTCATTGTCGCCACGC-3’) and FISH microscopy performed on dissected tissues using *Sod1238R* probe (3’-Cy3-TCCGCTGACTCTCGGGAGAT-5’). Indirect evidence for its vertical transmission came from the presence of this symbiont in the testes as well as the host’s egg surface and interior. The type strain, Nvir^T^, was isolated from salivary glands of *N. viridula* collected in 2019 feeding on host plants (black mustard, black nightshade, crown vetch, soybean seeds, flat beans). The draft genome size was 5 497 650 bp and the G+C content 55.6 mol%. The draft genome sequence of this strain has been deposited in the GenBank database under the Project number PRJEB70466 and submission ERA27452063. A previous assembly of this symbiont from a metagenomic sequencing effort of *N. viridula* can be found under accession number CAUIKD000000000. Further characterization of *S. nezarae*, as well as the deposition at the public repository will be described in a separate article.

## Discussion

*N. viridula* is a piercing and sucking insect that feeds on nutritionally imbalanced phloem sap, thus it relies on symbiotic associations with bacteria to biosynthesize essential nutrients (Tada et al., 2011). Pentatomidae shield bugs are associated with obligate *Pantoea* symbionts which colonize crypts in the M4 section of the gut (Duron & Noel, 2016), but little is known about other members of the shield bug microbial community. Hosokawa et al. (2015) suggested that *Sodalis* symbionts are facultatively associated with shield bugs, however, recent studies proposed an obligate nature of *Sodalis* in *N. viridula* due to its evident benefit for the host and a vertical transmission route (Fourie et al., 2023; Geerinck et al., 2022). *Sodalis* dominates the external egg-associated microbial community and, together with *Pantoea*, was highly abundant inside eggshells (Geerinck et al., 2022). This suggests that *Sodalis* is important to the host and most likely fulfils essential symbiotic functions to *N. viridula*. Along with that, the insect was shown to transmit *Sodalis* via saliva to plants, which repressed the biosynthesis of secondary plant metabolites allowing the insect to cope with plant defences (Coolen et al., 2024).

In this study, we comprehensively characterized the prevalence of the previously understudied *Sodalis* symbiont of laboratory-reared *N. viridula* in eggs, insect tissues and organs. Our findings revealed that *Sodalis* constituted an integral component of both the external and internal egg microbiome, confirming previous reports (Geerinck et al., 2022). The authors compared egg-associated microbiomes from *N. viridula* eggs obtained from two distinct geographical locations. Although they used a more stringent protocol to remove external egg microbiota in comparison to this study, their data revealed that their Belgian population had two primary egg-associated symbionts, *Pantoea* and *Sodalis.* In that study, *Sodalis* exceeded 70% of the relative abundance on the egg surface and 20% inside eggs. However, *Sodalis* was not found to be a part of the internal or external egg-associated microbiome in their Italian population. These variations, however, might be attributed to the difference in the insect subspecies as previously reported in other distinct populations of *N. viridula* (Geerinck et al., 2022; Hosokawa et al., 2016; Medina et al., 2018).

The eggs of the Pentatomidae family species are typically covered with a protective layer which is secreted by the females during oviposition (Shan et al., 2021). It forms a physical barrier for the developing nymphs preventing dehydration and pathogen entrance as well as protecting symbionts and secures their transmission to offspring. Prado et al. (2006) and Tada et al. (2011) found that surface sterilisation drastically decreased *N. viridula* vitality and development. However, our observations indicate that despite egg-surface sterilization, *Sodalis* and *Pantoea* were acquired in the adult *N. viridula* microbiome as evidenced by their presence in the microbial community profiles of the intestinal tract. This implies a likely origin of symbionts from the inside of the egg and a much more intimate relation with the host than symbionts transmitted through the egg surface. Nevertheless, the removal of symbionts from eggshells negatively impacted *N. viridula* survival and appearance. Surface sterilization decreased the abundance of *Sodalis* and *Pantoea*, and thus could result in less efficient colonization of the host and a decreased survival rate. Moreover, it led to the acquisition of *Enterobacter* and *Enterococcus* with simultaneous decrease in the relative abundance of *Sodalis*, suggesting potential repression of *Sodalis* growth by environmental microbes. The disturbance of the egg microbiota might have contributed to the easier invasion of entomopathogenic bacteria, which has adverse effects on insect fitness and survival rate (Tozlu et al., 2019). Shield bugs have a susceptible time window during the 2^nd^ instar period for the acquisition of symbionts, including those which are horizontally transmitted, explaining the presence of *Enterococcus* and *Enterobacter* in sterilized populations (Kikuchi et al., 2011). Although the efficiency of surface sterilization was not analysed in this study and others have used various methods to remove the external egg microbiome, our results demonstrate that factors such as environmental conditions and genetic traits could influence the performance and dynamics of shield bug populations. Furthermore, our findings strongly suggest the ability of *N. viridula* to retain *Sodalis* and *Pantoea* in the event of external egg microbiome disturbance and maintain vertical transmission of both microbes via internal storage of symbionts in eggs. Taken together, this underlines the complexity behind the manipulation of the egg microbiome, symbiont acquisition, and its effect on insect fitness, and therefore should be considered in the context of pest insect management strategies.

Through our investigation, we determined that the *Sodalis* symbiont is present in the majority of *N. viridula* organs, but predominant in the salivary glands, testes, and anterior regions of the midgut.

*Sodalis* symbionts were first reported in tsetse flies but have since been observed in weevils, psyllids, aphids, mealybugs, and various shield bugs (Ghosh et al., 2020; Hosokawa et al., 2015; Kaiwa et al., 2011; Koga & Moran, 2014). Although *Sodalis* infections are rare in most shield bug species, a high abundance of symbiont in *N. viridula* indicates either a resilience of *Sodalis*, similar to the one observed in insect-associated *Wolbachia* (Souto-Maior et al., 2015), or significant selective pressure favouring its presence (Hosokawa et al., 2015; Kaiwa et al., 2010, 2011). Furthermore, the structural organization of symbionts within specific organs is possibly controlled by the host. In honey bees, the colonization of microbes is orchestrated by the insect which secretes organic acids into the gut lumen favouring the growth of symbiotic *Snodgrassella alvi* (Quinn et al., 2024). On the other hand, Kim et al. (2013) described that insect midgut epithelia could produce antimicrobial substances and, in that way, control the selective infection of symbionts to the M4 midgut crypts. Moreover, the recent discovery of the sorting organ, a constricted region between the M3 and M4 section of the midgut, underscores its pivotal role in facilitating the shield bug gut symbiosis (Ohbayashi et al., 2015). Although it is unclear whether a comparable organ exists in *N. viridula*, a similar pattern was observed in the decreasing abundance of *Sodalis* along the intestinal tract. The analysis showed that *Sodalis* colonized anterior parts of the gut, including the M3 region, and no *Sodalis* was observed during imaging of the posterior M4 region including the crypts suggesting the presence of a sorting mechanism in *N. viridula*. The structural arrangement of *Sodalis* in specific organs is possibly linked to its function for the host. Hence, the colonization of salivary glands may be linked to *Sodalis’* ability to repress plant defences, while participation in digestion, detoxification, and nutrient supplementation (namely thiamine) could be associated with gut colonization (Coolen et al., 2024). Interestingly, the presence of Gammaproteobacteria and *Sodalis* in the testes and the absence in ovaries and unlaid eggs collected from the ovaries shown by others (Geerinck et al., 2022; Kikuchi et al., 2009), suggests that *N. viridula* males play a role in vertical symbiont transmission to the offspring. Tsetse flies maternally and paternally transmit obligate intracellular *Wolbachia* symbionts, illustrated by the detection of microbes in ovaries and testes (Doudoumis et al., 2012). Likewise, *S. glossinidius* was shown to be transmitted from males to females during mating (De Vooght et al., 2015) and Watanabe et al. (2014) discovered that the bacterial symbiont *Rickettsia* is vertically transmitted via sperm in the leafhopper *Nephotettix cincticeps*. Paternal transmission of bacteria was observed in *Anopheles stephensi* mosquitos too (Damiani et al., 2008). However, to date, there has been limited focus on the male reproductive organs within the Pentatomidae family. Whether parental transmission of symbionts via the internal egg microbiome occurs in *N. viridula* and other shield bugs is of interest and deserves future study.

Genomic analysis of *S. nezarae* revealed its close evolutionary relationship to *S. pierantonii* and *S. praecaptivus*, the latter representing a non-host-restricted lineage. Therefore, it served as a valuable reference of the ancestral state to infer *Sodalis* genome evolution (Clayton et al., 2012; Lo et al., 2016).

*S. praecaptivus* was described as an opportunistic human pathogen (Oakeson et al., 2014). Unlike other insect-associated microbes, a phylogenomic analysis showed no evidence of co-speciation events among *Sodalis* symbionts (Hosokawa et al., 2006; Kikuchi et al., 2009; Renoz et al., 2023), which raised questions regarding *Sodalis* acquisition by insects. Our comparative genomic analyses of *Sodalis* revealed that *S. nezarae* shared characteristics with both the free-living *S. praecaptivus* and the obligate host-associated *S. pierantonii*. *S*. *nezarae* displayed a unique large number of plasmids which suggests genome instability and adaptation towards a symbiotic lifestyle (Campbell et al., 2017; Van Leuven et al., 2014). Similar to *S. endolongispinus*, an obligate mealybug-associated *Sodalis* species, *S. nezarae* retained a genome containing thousands of pseudogenes, despite a similar genome size as HS1, indicating its recent shift from a free-living to an endosymbiotic lifestyle (Garber et al., 2021). In the context of insect symbiosis, *S. nezarae* probably fulfils diverse functions for the insect host, contributing to repression of plant defences, while participation in digestion, detoxification, and nutrient supplementation (Coolen et al., 2024; Manzano-Marín et al., 2023; Manzano-Marin et al., 2017; Renoz et al., 2023; Sloan & Ligoxygakis, 2017). Our findings point to the divergent nature of *S. nezarae* and show the strain’s likely recent evolutionary journey from a once free-living ancestor to becoming a vertically transmitted beneficial symbiont with a possible obligate role in *N. viridula*.

In conclusion, our research demonstrated the vertical transmission of both *Sodalis* and *Pantoea* symbionts within its eggs. Our findings suggest a potential role of males in this transmission process, as illustrated by the presence of bacteria within the testes microbiome. Besides this, the study provides evidence for selective control in the gut colonization orchestrated by *N. viridula* and reveals that previously overlooked *S. nezarae* is possibly an obligate symbiont that has undergone genome degeneration following an adaptation to symbiotic life. Altogether, our results present an example of the intimate relationship between insects and microbes which could be essential in the development of targeted pest control strategies in the future.

## Acknowledgements

We are thankful to the Radboud University Plant Ecology and Physiology departments for allowing us to make use of their insect cage growing facilities in the greenhouse. This study was supported by the Netherlands Organization for Scientific Research through the Gravitation Grant Netherlands Earth System Science Centre (grant number 024.002.001) and the Gravitation Grant Soehngen Institute of Anaerobic Microbiology (grant number 024.002.002) as well as the Radboud Institute for Biological Research (RIBES) and the Faculty of Science at Radboud University. The authors thank Prof. Dr Thomas Rattei and his team for maintaining the *Life Science Compute Cluster* (LiSC; https://cube.univie.ac.at/lisc) that was used for computational analyses.

## Competing Interests

The authors declare no competing interests.

## Data Availability Statement

Sequencing data are deposited in the European Nucleotide Archive under Project Number PRJEB70466 and submission ERA27452063. Supplementary material can be found in zenodo repository https://doi.org/10.5281/zenodo.10715847.

## Notes

https://doi.org/10.5281/zenodo.10715847

### Competing Interest Statement

The authors have declared no competing interest.

