## Supplementary Figures for "From Eggs to Guts: Symbiotic Association of *Sodalis nezarae* sp. nov. with the Southern Green Shield Bug *Nezara viridula*"

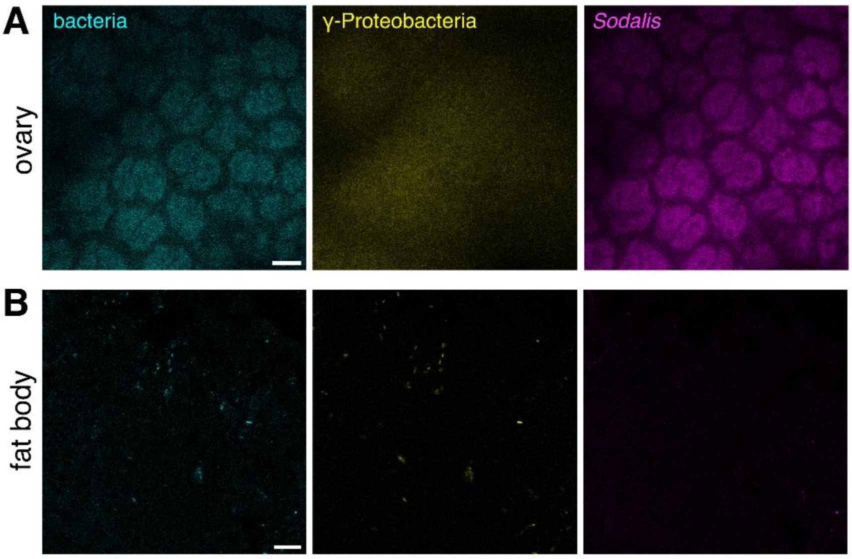


**Supplementary Figure 1. Colonization of *Nezara viridula* ovaries and fat body surface by Gammaproteobacteria and *Sodalis.*** Confocal micrographs show **A)** the adult *N. viridula* ovary, scale bar = 10 µm and **B)** the surface of the fat body, scale bar = 10 µm. Panel *bacteria* shows FISH-probe *Eub-mix* in cyan (Fluos). Panel *γ-Proteobacteria* shows FISH-probe *GAM42A* in yellow (Cy5). Panel *Sodalis* shows FISH-probe *Sod1238R* in magenta (Cy3). The images shown here are representative of FISH micrographs collected from three female insects.

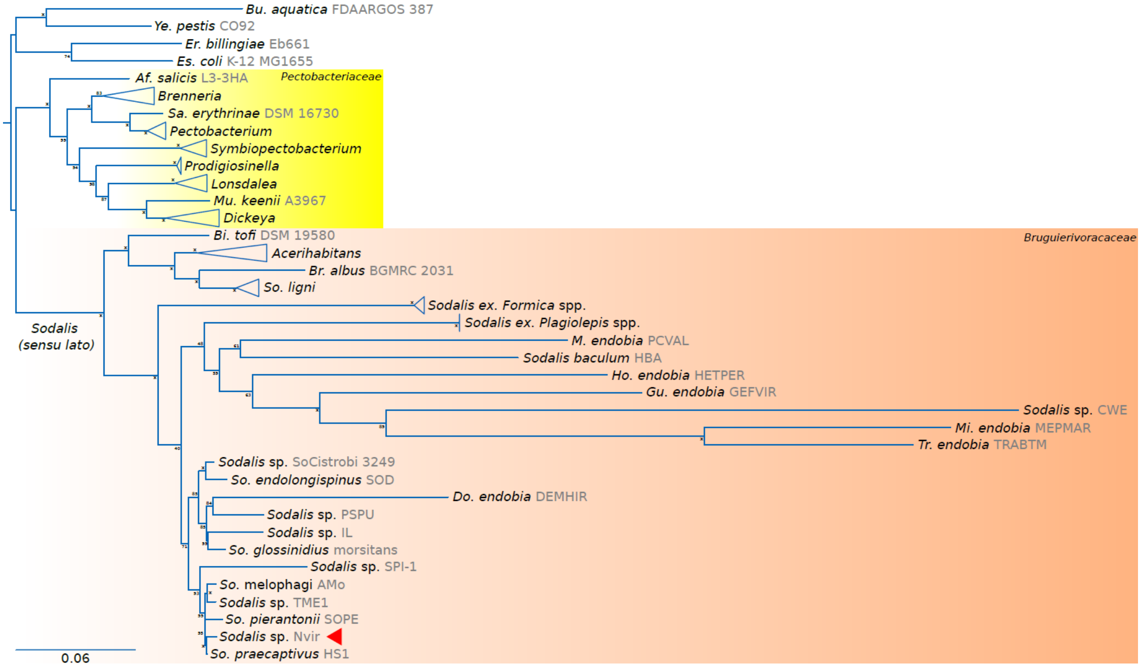


**Supplementary Figure 2. *Phylogenetic* relationships of *Pectobacteriaceae* and *Bruguierivoracaceae* genera.** Maximum-Likelihood phylogenetic tree as calculated using IQ-TREE v2.2.2.7 (LG4X+I+G; 1000 UltraFast bootstrap replicates). Names on leaves specify the bacterial genera, species, and strains (in grey). Coloured boxes delimit the *Pectobacteriaceae* and the proposed "*Bruguierivoracaceae*" (or *Sodalis sensu lato*). A red left-pointing arrow highlights the novel *Sodalis* sp. Nvir. Values at nodes indicate the UltraFast bootstrap support values in percentages. An asterisk (*) denotes a support of 100%.

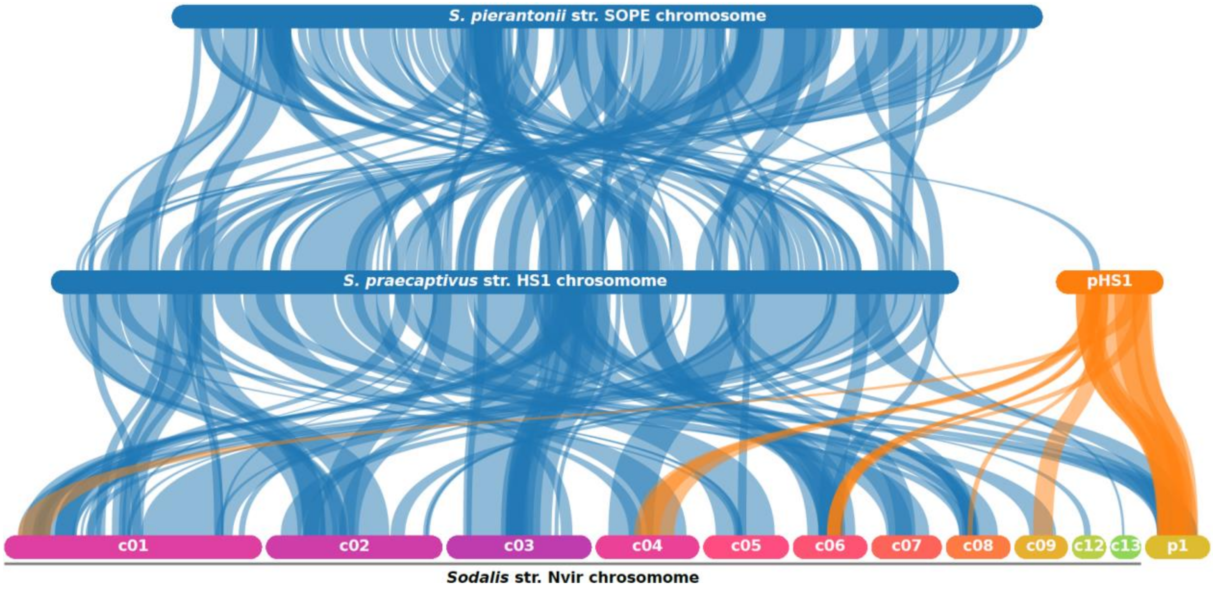


**Supplementary Figure 3. Syntenic blocks among selected *Sodalis* strains.** Coloured horizontal bars represent chromosomes/contigs, while vertical curved lines connect syntenic blocks. The latter are colour-coded in relation to the former. A "c" and "p" prefix in horizontal bar labels marks the molecule as a contig or a plasmid molecule, respectively. For simplicity, only molecules that share syntenic blocks are displayed. Horizontal bars are at scale in relation to the size of the molecule (in bps).
