## Supplementary Methods for "From Eggs to Guts: Symbiotic Association of *Sodalis nezarae* sp. nov. with the Southern Green Shield Bug *Nezara viridula*"

**Supplementary Information Methods**

**Genomic annotation and metabolic pathway prediction**

To annotate the genes involved in the biosynthesis of essential amino acids, B vitamins, and selected co-factors in *Sodalis* sp. Nvir, we collected the sequences of the proteins involved in these pathways from *S. praecaptivus* HS1. In addition, we collected the same set of proteins from *Escherichia coli* K-12 MG1655 to annotate those genes in the previously sequenced primary obligate *Pantoea* symbiont of *N. viridula* (INSDC: CAUIKC010000001.1, OV793556.1). Guide pathways were extracted from EcoCyc (Keseler et al., 2021).

For phylogenetic placement of *Sodalis* strains, collected genomes were submitted to OrthoFinder v2.5.4 (Emms & Kelly, 2019) to group predicted protein-coding genes across selected genomes in families of orthologous proteins. Then, we aligned all single-copy marker genes (i.e. present across all genomes) using MAFFT v7.490 (-maxiterate 1000 -localpair; (Katoh & Standley, 2013)), concatenated them, and then subjected the alignment to Gblocks v0.91b (Talavera & Castresana, 2007) for removal of divergent and ambiguously aligned blocks.

**ANI and 16S rRNA gene identity scoring**

To calculate pairwise ANI scores, we extracted the genomic sequences for selected relatives of *Sodalis* sp. Nvir. These sequences were used as input for OthoANIu v1.2 (Lee et al., 2016). For guidance, species and genus thresholds were set following Chun et al. (2018) and Barco et al. (2020).

For calculating 16S rRNA gene identity, selected 16S rRNA gene sequences were extracted and aligned using the built-in Muscle v3.8.425 (Edgar, 2021) from AliView v1.28 (Larsson, 2014). Last, Clustal Omega v1.2.4 (Sievers et al., 2011) was used to calculate a pairwise 16S rRNA gene identity matrix. The species and genus level threshold were set following Kim et al. (2014) and Yarza et al. (2014).

**Phage region identification**

To infer prophage regions across the selected *Sodalis* genomes, we extracted the nucleotide sequences for all molecules (*e.g.* chromosomes, plasmids, contigs) per organism. These were then used as input for PHASTEST (Arndt et al., 2016; Wishart et al., 2023; Zhou et al., 2011), as implemented in the web server (https://phastest.ca; last accessed 15^th^ February 2024). Visualisation and export of diagrams were done using the built-in visualisation tool.

**Diagnostic PCR**

The presence of *Sodalis* and *Pantoea* symbionts in sterilized and control egg masses was determined with a PCR reaction quantifying *gro*L gene using specific primers *SodF* (5’- CCCTTATCGATAGCCGCGTT-3’) and *SodR* (5’-GATCTTCATTGTCGCCACGC-3’) and *PantF* (5’-TCGCAGTTCTAAAGGTGGGT-3’) and *PantR* (5’-GTAGCGCCACTTTAATGCCA-3’), respectively (Coolen, Rogowska-van der Molen et al. *under review*). Additionally, *Sodalis* presence in organs of adult *N. viridula* male and female was determined using the same *Sodalis*-specific primer pair. Amplification was performed using SensoQuest lab-cycler (BIOKÉ, the Netherlands) in a 25 mL reaction volume and consisted of 12.5 µL PerfeCTa Quanta SYBR Green FastMix (Quanta Bio, USA), forward and reverse primers of the final concentration of 0.1 pmol µL^-1^, 1 μL of template DNA and 9.5 μL Nuclease Free Water (Thermo Fisher Scientific). The PCR programme was initiated by initial denaturation at 95 °C for 3 min, followed by 30 cycles of denaturation at 95 °C for 30 s, annealing at 59 °C for 30 s, elongation at 72 °C for 30 s, and 5 min at 72 °C for the final elongation. PCR products were examined by electrophoresis at 80 V for 1 h in a 1.5% (w/v) agarose gel containing ethidium bromide in 1 x SB buffer. To determine PCR product size GeneRuler 100 bp DNA ladder was used as a marker DNA (Thermo Fisher Scientific). The electrophoresis gel was examined under UV light.

**Fluorescence *in situ* hybridization**

To image total bacteria, Gammaproteobacteria and the *Sodalis*, we dissected the fat body, ovary, testes, salivary glands, M1, M2, M3, M4 sections of the midgut, hindgut and Malpighian tubules from one adult male and one female *N. viridula* as described under *Insect dissection*. Isolated organs were washed twice with PBS and incubated with 300 µL PBS and 900 µL paraformaldehyde solution (4% paraformaldehyde in PBS) for 2 h on ice and samples were washed twice with 500 µL PBS. The fixed tissue samples were placed on SuperFrost Plus™ adhesion microscope slides (Epredia™, USA) and dehydrated in increasing ethanol concentration (50%, 80%, 100%) for 10 min each. Then the samples were air-dried and incubated for 3 h at 46 °C with a hybridization buffer (0.9 M NaCl, 0.02M Tris-HCl (pH 8.0), 35% formamide, 0.02% sodium dodecyl sulfate) and specific probes in the humidified chamber. For total bacterial detection and identification of specific bacterial groups within the samples, fluorescence *in situ* hybridization (FISH) using a Fluos-labelled general bacterial probe (Eub-mix; (Amann et al., 1990; Daims et al., 1999)) and a Cy5-labeled Gammaproteobacterial probe (GAM42A; (Manz et al., 1992)) along with a GAM42A competitor probe were employed. Along, a Cy3-labeled probe specific for *Sodalis* sp. was used for targeted detection (Sod1238R; (Koga et al., 2013)). The final concentration of Cy3 and Cy5-probes in the hybridization buffer was 0.5 pmol µL^-1^ (Sod1238R and GAM42A) and a Fluos-probe 0.83 pmol µL^-1^ (Eub-mix).

After hybridization, tissue samples were washed in the wash buffer (0.2 M Tris-HCl (pH 8), 0.08 M NaCl, 5 mM EDTA) for 10 min at 48 °C, dipped in ice-chilled ultrapure Milli-Q® water, air-dried and embedded in Vectashield without 4,6-diamidino-2-phenylindole (Vector Laboratories, Burlingame, CA, USA). Images were taken using a confocal laser scanning microscope (Leica SP8X, Mannheim, Germany) and image processing was performed in Fiji (Schindelin et al., 2012).

Amann, R. I., Binder, B. J., Olson, R. J., Chisholm, S. W., Devereux, R., & Stahl, D. A. (1990). Combination of 16S rRNA-targeted oligonucleotide probes with flow cytometry for analyzing mixed microbial populations. *Appl Environ Microbiol*, *56*(6), 1919-1925. <https://doi.org/10.1128/aem.56.6.1919-1925.1990>

Arndt, D., Grant, J. R., Marcu, A., Sajed, T., Pon, A., Liang, Y., & Wishart, D. S. (2016). PHASTER: a better, faster version of the PHAST phage search tool. *Nucleic Acids Res*, *44*(W1), W16-21. <https://doi.org/10.1093/nar/gkw387>

Barco, R. A., Garrity, G. M., Scott, J. J., Amend, J. P., Nealson, K. H., & Emerson, D. (2020). A Genus Definition for Bacteria and Archaea Based on a Standard Genome Relatedness Index. *mBio*, *11*(1). <https://doi.org/10.1128/mBio.02475-19>

Chun, J., Oren, A., Ventosa, A., Christensen, H., Arahal, D. R., da Costa, M. S., Rooney, A. P., Yi, H., Xu, X. W., De Meyer, S., & Trujillo, M. E. (2018). Proposed minimal standards for the use of genome data for the taxonomy of prokaryotes. *Int J Syst Evol Microbiol*, *68*(1), 461-466. <https://doi.org/10.1099/ijsem.0.002516>

Daims, H., Brühl, A., Amann, R., Schleifer, K. H., & Wagner, M. (1999). The domain-specific probe EUB338 is insufficient for the detection of all Bacteria: development and evaluation of a more comprehensive probe set. *Syst Appl Microbiol*, *22*(3), 434-444. <https://doi.org/10.1016/s0723-2020(99)80053-8>

Edgar, R. C. (2021). MUSCLE v5 enables improved estimates of phylogenetic tree confidence by ensemble bootstrapping. *bioRxiv*, 2021.2006.2020.449169. <https://doi.org/10.1101/2021.06.20.449169>

Emms, D. M., & Kelly, S. (2019). OrthoFinder: phylogenetic orthology inference for comparative genomics. *Genome Biol*, *20*(1), 238. <https://doi.org/10.1186/s13059-019-1832-y>

Katoh, K., & Standley, D. M. (2013). MAFFT multiple sequence alignment software version 7: improvements in performance and usability. *Mol Biol Evol*, *30*(4), 772-780. <https://doi.org/10.1093/molbev/mst010>

Keseler, I. M., Gama-Castro, S., Mackie, A., Billington, R., Bonavides-Martinez, C., Caspi, R., Kothari, A., Krummenacker, M., Midford, P. E., Muniz-Rascado, L., Ong, W. K., Paley, S., Santos-Zavaleta, A., Subhraveti, P., Tierrafria, V. H., Wolfe, A. J., Collado-Vides, J., Paulsen, I. T., & Karp, P. D. (2021). The EcoCyc Database in 2021. *Front Microbiol*, *12*, 711077. <https://doi.org/10.3389/fmicb.2021.711077>

Kim, M., Oh, H. S., Park, S. C., & Chun, J. (2014). Towards a taxonomic coherence between average nucleotide identity and 16S rRNA gene sequence similarity for species demarcation of prokaryotes. *Int J Syst Evol Microbiol*, *64*(Pt 2), 346-351. <https://doi.org/10.1099/ijs.0.059774-0>

Koga, R., Bennett, G. M., Cryan, J. R., & Moran, N. A. (2013). Evolutionary replacement of obligate symbionts in an ancient and diverse insect lineage. *Environ Microbiol*, *15*(7), 2073-2081. <https://doi.org/10.1111/1462-2920.12121>

Larsson, A. (2014). AliView: a fast and lightweight alignment viewer and editor for large datasets. *Bioinformatics*, *30*(22), 3276-3278. <https://doi.org/10.1093/bioinformatics/btu531>

Lee, I., Ouk Kim, Y., Park, S. C., & Chun, J. (2016). OrthoANI: An improved algorithm and software for calculating average nucleotide identity. *Int J Syst Evol Microbiol*, *66*(2), 1100-1103. <https://doi.org/10.1099/ijsem.0.000760>

Manz, W., Amann, R., Ludwig, W., Wagner, M., & Schleifer, K.-H. (1992). Phylogenetic Oligodeoxynucleotide Probes for the Major Subclasses of Proteobacteria: Problems and Solutions. *Systematic and Applied Microbiology*, *15*(4), 593-600. <https://doi.org/https://doi.org/10.1016/S0723-2020(11)80121-9>

Schindelin, J., Arganda-Carreras, I., Frise, E., Kaynig, V., Longair, M., Pietzsch, T., Preibisch, S., Rueden, C., Saalfeld, S., Schmid, B., Tinevez, J. Y., White, D. J., Hartenstein, V., Eliceiri, K., Tomancak, P., & Cardona, A. (2012). Fiji: an open-source platform for biological-image analysis. *Nat Methods*, *9*(7), 676-682. <https://doi.org/10.1038/nmeth.2019>

Sievers, F., Wilm, A., Dineen, D., Gibson, T. J., Karplus, K., Li, W., Lopez, R., McWilliam, H., Remmert, M., Soding, J., Thompson, J. D., & Higgins, D. G. (2011). Fast, scalable generation of high-quality protein multiple sequence alignments using Clustal Omega. *Mol Syst Biol*, *7*, 539. <https://doi.org/10.1038/msb.2011.75>

Talavera, G., & Castresana, J. (2007). Improvement of phylogenies after removing divergent and ambiguously aligned blocks from protein sequence alignments. *Syst Biol*, *56*(4), 564-577. <https://doi.org/10.1080/10635150701472164>

Wishart, D. S., Han, S., Saha, S., Oler, E., Peters, H., Grant, J. R., Stothard, P., & Gautam, V. (2023). PHASTEST: faster than PHASTER, better than PHAST. *Nucleic Acids Res*, *51*(W1), W443-W450. <https://doi.org/10.1093/nar/gkad382>

Yarza, P., Yilmaz, P., Pruesse, E., Glockner, F. O., Ludwig, W., Schleifer, K. H., Whitman, W. B., Euzeby, J., Amann, R., & Rossello-Mora, R. (2014). Uniting the classification of cultured and uncultured bacteria and archaea using 16S rRNA gene sequences. *Nat Rev Microbiol*, *12*(9), 635-645. <https://doi.org/10.1038/nrmicro3330>

Zhou, Y., Liang, Y., Lynch, K. H., Dennis, J. J., & Wishart, D. S. (2011). PHAST: a fast phage search tool. *Nucleic Acids Res*, *39*(Web Server issue), W347-352. <https://doi.org/10.1093/nar/gkr485>
